# Oxidative reactions catalyzed by hydrogen peroxide produced by *Streptococcus pneumoniae* and other Streptococci Cause the Release and Degradation of Heme from Hemoglobin

**DOI:** 10.1101/2022.08.23.504964

**Authors:** Babek Alibayov, Anna Scasny, Faidad Khan, Aidan Creel, Perriann Smith, Ana G. Jop Vidal, Fa’alataitaua M. Fitisemanu, Teresita Padilla-Benavides, Jeffrey Weiser, Jorge E. Vidal

## Abstract

*Streptococcus pneumoniae* (Spn) strains cause pneumonia that kills millions every year worldwide. Spn produces Ply, a hemolysin that lyses erythrocytes releasing hemoglobin and also produces the pro-oxidant hydrogen peroxide (Spn-H_2_O_2_) during growth. The hallmark of the pathophysiology of hemolytic diseases is the oxidation of hemoglobin but oxidative reactions catalyzed by Spn-H_2_O_2_ has been poorly studied. We characterized the oxidation of hemoglobin by Spn-H_2_O_2_. We prepared a series of single (Δ*spxB*, or Δ*lctO*), double mutant (Δ*spxB*Δ*lctO*) and complemented strains in TIGR4, D39 and EF3030. We then utilized an *in vitro* model with oxy-hemoglobin to demonstrate that oxy-hemoglobin was oxidized rapidly, within 30 min of incubation, by Spn-H_2_O_2_ to met-hemoglobin and that the main source of Spn-H_2_O_2_ was pyruvate oxidase (SpxB). Moreover, extended incubation caused the release and the degradation of heme. We then assessed oxidation of hemoglobin and heme degradation by other bacteria inhabitants of the respiratory tract. All hydrogen peroxide-producing streptococci tested caused the oxidation of hemoglobin and heme degradation whereas those bacterial species that produce <1 μM H_2_O_2_, neither oxidized hemoglobin nor degraded heme. An *ex vivo* bacteremia model confirmed that oxidation of hemoglobin and heme degradation occurred concurrently with hemoglobin that was released from erythrocytes by Ply. Finally, gene expression studies demonstrated that heme, but not red blood cells or hemoglobin induced an upregulated transcription of the *spxB* gene. Oxidation of hemoglobin may be important for pathogenesis and for the symbiosis of hydrogen peroxide-producing bacteria with other species by providing nutrients such as iron.

## Introduction

*Streptococcus pneumoniae* (Spn) colonizes the lower respiratory epithelium, causing pneumonia that can progress to invasive pneumococcal disease (IPD)(1–4). Spn mainly affects the most vulnerable populations: young children, immunocompromised patients, and the elderly (1, 4, 5). Globally, Spn causes ~15 million cases of illness each year, leading to more than a million deaths due to pneumonia (6–8). Pneumonia is an acute infection of the pulmonary parenchyma that presents with pulmonary hemorrhage, inflammatory congestion, hepatization, suppurative infiltration and lung parenchymal injury (4, 9–13). If left untreated, Spn invades the bloodstream, producing septic shock, multiorgan failure, cardiotoxicity, and death within days from onset (4, 9–12, 14, 15).

Spn strains produce the toxin pneumolysin (Ply) that has cytotoxic activity against lung epithelial and endothelial cells (16–18). Ply is a 53-kDa surface located protein (19, 20) that can also be released through autolysis (21, 22). Besides its toxicity against cells, Ply has hemolytic activity against erythrocytes of several mammalian species and hemolysis is observed whether Ply is released into the supernatant or located on the bacterial membrane (20). Under standard culture conditions, i.e., growing bacteria at 37°C in a 5% CO_2_ atmosphere, Ply is responsible for almost all hemolytic activity in cultures of Spn strains (11), causing the release of a large amount of hemoglobin into the supernatant (20, 21, 23). Transcription of the *ply* gene encoding this pneumolysin is upregulated when Spn is cultured under conditions mimicking lung infection (24) and therefore, Ply-induced release of hemoglobin is expected to occur during pneumococcal lung infection.

Hemoglobin is a tetrameric protein (64 kDa) made of four subunits, two alpha and two beta chains. Each subunit contains a heme center that reversibly binds oxygen through a penta-coordinate heme molecule containing ferrous iron (Fe^+2^), known as oxy-hemoglobin (25, 26). When hemoglobin is released from erythrocytes, heme-hemoglobin can be observed by spectroscopy at ~415 nm (25, 27, 28). This region is known as the Soret region peak and represents heme-hemoglobin, while the alpha and beta chains are characterized by two absorption peaks of ~540 and ~570 nm (27, 28). Cell-free heme is toxic to cells because of its potential to generate toxic radicals (29).

Release of hemoglobin during hemolytic disease such as sickle cell disease and other hemorrhagic conditions, causes oxidation of hemoglobin (30). Oxidized hemoglobin releases heme, which in turn induces adverse effects including leukocyte activation, cytokine up-regulation, and production of oxidants leading to lung epithelial injury and damage to the vascular system (29–31). Reactive oxygen species (ROS), such as nitric oxide (NO) (32) hypochlorous acid conjugate base (OCl-)(24), and H_2_O_2_ (33, 34), drive a catalytic cycle including the oxidation of oxy-hemoglobin to ferryl hemoglobin (Hb-Fe^+4^), and auto - reduction of the ferryl intermediate to met-hemoglobin (Hb-Fe^+3^) that is detected by spectroscopy at 405 nm (35, 36). Additional pro-oxidant molecules, such as H_2_O_2_, produce more ferryl hemoglobin and cause heme degradation (28, 36). Thus, the oxidation of hemoglobin results in highly oxidizing ferryl hemoglobin (Hb-Fe^4+^), met-hemoglobin (Hb-Fe^3+^) and the release of toxic heme that disrupts the mitochondria membrane potential of lung epithelial cells and endothelial cells, and caused lipid peroxidation (28, 30, 36–38).

Because Spn causes hemorrhage in the lung and produces a large amount of highly reactive H_2_O_2_ (39, 40), oxidation of cell-free hemoglobin might occur during pneumococcal disease. H_2_O_2_ is a by-product of the oxidation of pyruvate, end product of the glycolysis, produced by the enzyme pyruvate oxidase (SpxB) (41, 42). This reaction consumes oxygen and produces H_2_O_2_. In the absence of oxygen, pyruvate is converted to L-lactate while in the presence of molecular O_2_, L-lactate feeds back to pyruvate, via lactate oxidase (LctO), in a reaction that also produces H_2_O_2_ (41, 42). It has been calculated that ~85% of H_2_O_2_ released in cultures of Spn occurs through the reaction catalyzed by SpxB, whereas the remaining ~15% is attributed to LctO (43, 44). Experiments using animal models have demonstrated that Spn-H_2_O_2_ plays a role in lung colonization, and in the translocation of Spn from the lungs to the bloodstream (45–48). Details of the mechanism(s) are largely unknown.

We recently discovered that Spn-H_2_O_2_ oxidizes hemoglobin to met-hemoglobin, a non-oxygen binding form of hemoglobin (23). We also demonstrated that the oxidation of hemoglobin causes the characteristic greenish halo around pneumococcal colonies grown on blood agar plates, known as alpha hemolysis (23). Thus, the phenotype known as alpha hemolysis is not hemolysis but the oxidation of hemoglobin to met-hemoglobin (23). Another alpha-hemolytic streptococci, *S. gordonii*, oxidizes hemoglobin to met-hemoglobin through catalytic reactions driven by H_2_O_2_ (49).

Oxidation of cell-free hemoglobin induces lung injury in patients with acute respiratory distress syndrome (31), and in children with SCD, oxidation of hemoglobin (Hb)S to Hb-Fe^4+^ and then its reduction to met-hemoglobin results in an acute hemolytic vascular inflammatory process causing acute lung injury (30, 50). We hypothesize that the oxidation of hemoglobin during pneumococcal disease can exacerbates cytotoxicity in the lungs but details of these oxidative reactions in cultures of Spn strains are not available. Therefore, in this study we comprehensively investigated the oxidation of hemoglobin by Spn cultures. We discovered that the oxidative capacity of Spn cultures, under the culture conditions utilized here, was attributed to H_2_O_2_ produced through catalytic reactions of SpxB but not to that of LctO. We also found that Spn-H_2_O_2_ caused the degradation of heme and that a number of H_2_O_2_-producing, alpha-hemolytic, Streptococci oxidized hemoglobin and degraded heme from hemoglobin. An *ex vivo* model of pneumococcal bacteremia further demonstrated that oxidation of hemoglobin and heme degradation occurred rapidly and that heme, but not hemoglobin or red blood cells, caused an upregulated transcription of *spxB*. Oxidative reactions driven by Spn-H_2_O_2_ might contribute to the pathogenesis of pneumococcal disease and can also be important for the symbiosis with other non-hydrogen peroxide producing bacteria by providing nutrients such as heme-iron.

## Results

### *Streptococcus pneumoniae*-produced hydrogen peroxide oxidizes heme-hemoglobin, causing the loss of the Soret peak

We demonstrated that the alpha and beta chains of oxy-hemoglobin are deoxygenated by *S. pneumoniae* strains but further oxidation of heme-Fe^+2^ was not investigated (23). To assess this, Todd-Hewitt broth (THY) supplemented with sheep oxy-hemoglobin (oxyHb), which contains heme-Fe^2+^, was inoculated with TIGR4 and incubated at 37°C in a 5% CO_2_ and ~20% O_2_ atmosphere. As a control, uninfected THY-oxyHb was incubated under the same conditions. After 30 min of incubation, TIGR4 cultures caused the shift of the Soret curve from heme-Fe^2+^, that peaks at 415 nm, to a Soret curve of heme-Fe^3+^ at 405 nm (Fig. 1A). As expected, this hemoglobin preparation was deoxygenated producing met-hemoglobin (Fig. 1B). Similar oxidation of heme-Fe^2+^to heme-Fe^3+^ was observed when oxy-hemoglobin was purified from horse erythrocytes and incubated with Spn strains TIGR4, or D39 (not shown). Cultures of THY-oxyHb inoculated with hydrogen peroxide-deficient strain TIGR4Δ*spxB*Δ*lctO*, or D39Δ*spxB*Δ*lctO*, did not cause the shift of the Soret curve (Fig. 1A and not shown). Hemoglobin in uninfected cultures was not auto-oxidized after this 30 min incubation period (Fig. 1A).

**Fig. 1.**
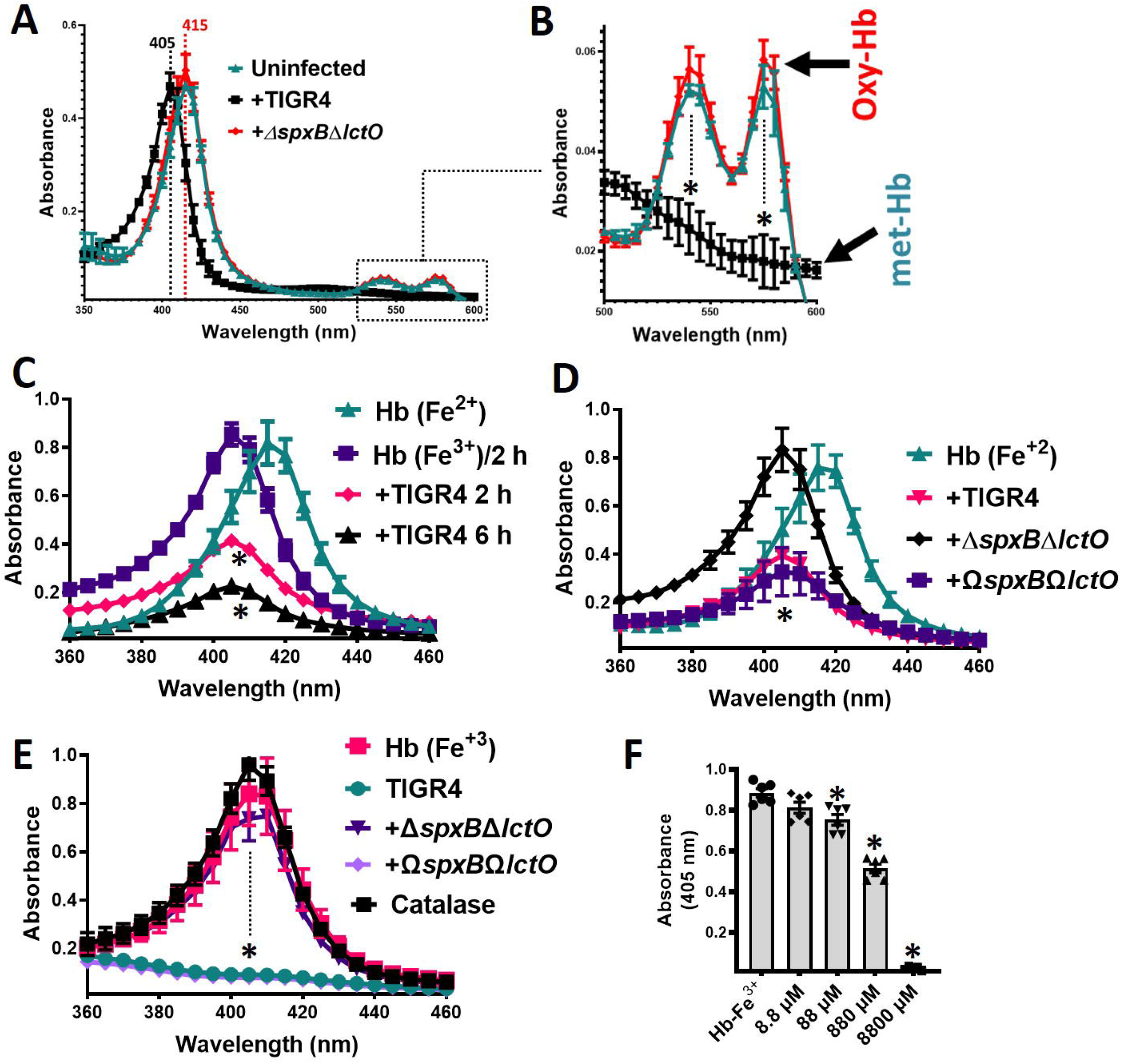
Hydrogen peroxide produced in *S. pneumoniae* cultures oxidizes human hemoglobin. TIGR4, TIGR4Δ*spxB*Δ*lctO*, or TIGR4Ω*spxB*Ω*lctO* was inoculated in THY containing oxy-hemoglobin [Hb (Fe^+2^)] or met-hemoglobin [Hb (Fe^+3^)], or left uninfected, and cultures were incubated for (A-B) 30 min, (C) 2 h or 6 h, (D-F) 6 h at 37°C in a 5% CO_2_ atmosphere. Supernatants were collected and the spectra were obtained using a spectrophotometer Omega BMG LabTech. Panel B shows the enclosed area in A, corresponding to the absorbance peaks of the alpha chain and the beta chain of oxy-hemoglobin, flattened in met-hemoglobin. In (A, B and E) the dotted lines indicate the Soret peak for oxy-hemoglobin (415 nm) or met-hemoglobin (405 nm). (F) THY supplemented with methemoglobin (Hb-Fe^3+^) was added with the indicated concentration of hydrogen peroxide, or left untreated. These cultures were incubated for 6 h at 37°C in a 5% CO_2_ atmosphere after which the supernatants were purified and the spectra were obtained as above. The absorbance at 405 nm obtained for each assessed concentration, was utilized to construct the graphic. Error bars in all panels represent the standard errors of the means calculated using data from at least three independent experiments performed with two technical replicates. In (B-F) One-way ANOVA with Dunnet’s test for multiple comparison was performed, \**p*<0.05 compared against the Soret peak of uninfected THY-Hb-Fe^2+^ (B and D), uninfected THY-Hb-Fe^2+^ and incubated 2 h (C) or uninfected THY-Hb-Fe^3+^(E and F).

Extended incubation of TIGR4 in THY-oxyHb revealed that the Soret curve was lost in a time-dependent manner (Fig. 1C). To assess if Spn-H_2_O_2_ had caused the loss of the Soret peak, we inoculated THY-oxyHb cultures with TIGR4Δ*spxB*Δ*lctO* and the culture was incubated for 6 h. A Soret curve was observed in this culture but the heme-Fe^2+^ peak (415 nm), observed after 30 min postinoculation (Fig. 1A), had shifted to 405 nm representing heme-Fe^3+^ (Fig. 1D). The deoxygenation of hemoglobin in THY-oxyHb cultures of TIGR4Δ*spxB*Δ*lctO* was, however, due to autoxidation because uninfected cultures incubated under the same conditions at 37°C, and with a 5% CO_2_ and ~20% O_2_ atmosphere, had shifted to heme-Fe^3+^ as well (Fig. 1C).

To further confirm that the loss of the Soret peak was due to Spn-H_2_O_2_, we prepared a complemented strain by cloning back *spxB* and *lctO* (Ω*spxB*Ω*lctO*) between SP_1113 and SP_1114 (Fig. 2). We also complemented TIGR4Δ*spxB* with a single copy of *spxB* to be utilized later (Fig. 2). This chromosomal location is unaffected by exogenous regulatory elements, such that complementing genes were controlled by their own promoters and insertion of genes within SP_1113 and SP_1114 did not affect virulence (51–54). Compared to wt and TIGR4Δ*spxB*Δ*lctO*, complemented strains showed a slight but not significant growth delay (Fig. 2B); however H_2_O_2_ production was restored to wt levels, as assessed with the Prussian blue-forming reaction and the Amplex Red kit (Fig. 2C and 2D) (55). In THY-oxyHb cultures of Ω*spxB*Ω*lctO* incubated for 6 h, the Soret curve had flattened (Fig. 1D). These results suggest that Spn-H_2_O_2_ causes heme degradation.

**Fig. 2.**
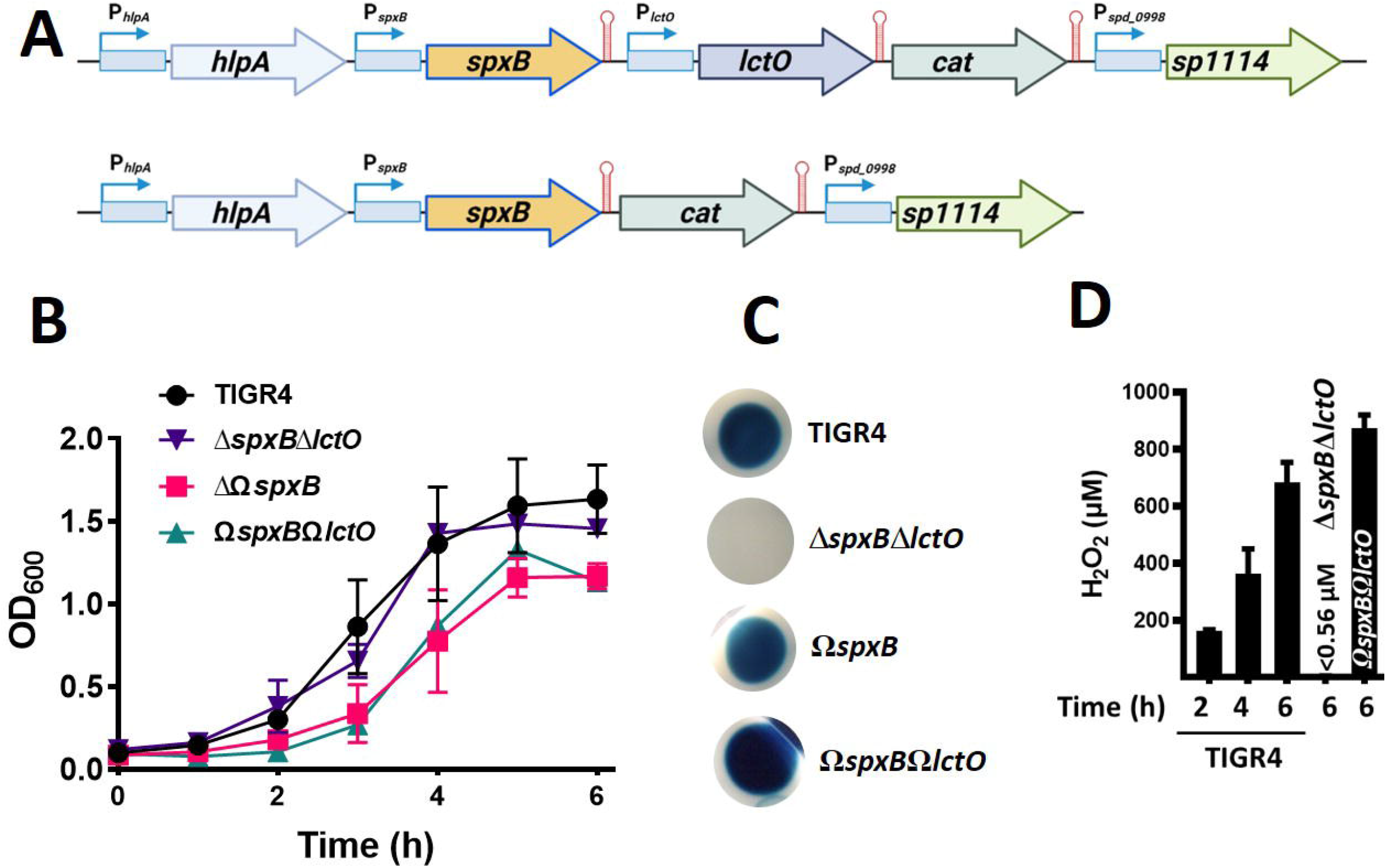
Preparation of a TIGR4Δ*spxB*Δ*lctO*-derivative complemented strain carrying *spxB* and *lctO* or *spxB*. (A) Schematic diagram showing the chromosomal region where *spxB* and *lctO* (top) or *spxB* (bottom) was complemented in TIGR4Δ*spxB*Δ*lctO*, to create TIGR4Ω*spxB*Ω*lctO*, TIGR4Ω*spxB*. Promotor regions of each gene (P) are indicated as well as terminators (†); gene IDs were included using TIGR4 nomenclature. (B-D) TIGR4, TIGR4Δ*spxB*Δ*lctO*, TIGR4Ω*spxB*Ω*lctO*, or TIGR4Ω*spxB* was inoculated in THY and incubated at 37°C in a 5% CO_2_ atmosphere. (B) The OD_600_ of the cultures was obtained at the indicated time. Error bars represent the standard errors of the means calculated using data from at least three independent experiments performed with two technical replicates. (C) Supernatants from 4 h cultures were sterilized and an aliquot was inoculated onto Prussian-blue agar plates (55) and the plates were incubated 2 h at room temperature after which they were photographed. This experiment was repeated at least three times. (D) Hydrogen peroxide was quantified from these supernatants using the Amplex Red assay. Error bars represent the standard errors of the means calculated using data from two independent experiments performed with two technical replicates. Limit of detection=0.56 μM.

Given that heme-Fe^2+^ in THY-oxyHb cultures was oxidized to heme-Fe^2+^ (Fig. 1A) prior to losing the Soret curve (Fig. 1C), we next assessed if the oxidation state of heme was important for the subsequent loss of the Soret curve. To investigate this, we supplemented THY with deoxyhemoglobin (i.e., met-hemoglobin) containing heme-Fe^3+^ (THY-Hb) and the culture was incubated with Spn. A similar loss of the Soret curve was observed with TIGR4 wt and Ω*spxB*Ω*lctO* (Fig. 1E), but an intact curve was obtained in cultures of TIGR4Δ*spxB*Δ*lctO* (Fig. 1E), indicating that heme-Fe^3+^ was further degraded. These results demonstrated that Spn-H_2_O_2_ converts oxyHb to metHb.

### Exogenous hydrogen peroxide supplementation at levels produced in *S. pneumoniae* cultures recapitulates hemoglobin oxidation

We then assessed whether a similar concentration of hydrogen peroxide to that observed in wt cultures of Spn (<1 mM) [Fig. 2D and references (39, 40)] can alter the heme-Fe^3+^ curve. For easy comparison of the multiple concentrations utilized in this experiment, we plotted the maximum absorbance obtained by spectroscopy for THY-Hb (405 nm) and we compared against that of THY-Hb treated with an increasing concentration of hydrogen peroxide. As shown in Fig. 1F, 10 μM hemoglobin in THY-Hb peaks at an absorbance of ~0.8. After 4 h of incubation with hydrogen peroxide, the absorbance was significantly reduced with a concentration of 88 μM and above, including the concentration of hydrogen peroxide observed in Spn cultures (~880 μM) (Fig. 1F). Altogether, these results indicate that the heme moiety in oxidized hemoglobin is degraded by Spn-H_2_O_2_.

### Oxidation of hemoglobin in *S. pneumoniae* cultures is mainly caused by hydrogen peroxide catalyzed through pyruvate oxidase SpxB

Under aerobic conditions, hydrogen peroxide in pneumococcal cultures is produced through catalytic reactions of pyruvate oxidase (SpxB) and lactate oxidase (LctO) (43, 44). To assess the contribution of each enzyme, single Δ*spxB* or Δ*lctO* prepared in the TIGR4, or EF3030, background were assessed in THY-Hb cultures. In cultures of wt strains TIGR4, and EF3030, isogenic mutant TIGR4Δ*lctO*, and EF3030Δ*lctO*, Ω*spxB*Ω*lctO* and TIGR4Δ*spxB*Ω*spxB* complemented strain the Soret Heme-Hb^3+^ curve was flattened, causing a statistically significant reduction of the 405 nm peak (Fig. 3A). In contrast, in THY-Hb cultures of TIGR4Δ*spxB* or EF3030Δ*spxB*, the Soret curve remained intact, similar to THY-Hb cultures inoculated with the corresponding Δ*spxB*Δ*lctO* mutant (Fig. 3A and 3B). Additional D39Δ*spxB*Δ*lctO*, or D39Δ*spxB* strain (P878), hydrogen peroxide-deficient mutants did not alter the Soret curve while D39Δ*spxB/spxB* complemented strain, P1221, caused the loss of the Soret curve (Fig. 3C). These experiments demonstrated that oxidation of hemoglobin in cultures of Spn is mainly caused by hydrogen peroxide sourced from reactions catalyzed by SpxB.

**Fig. 3.**
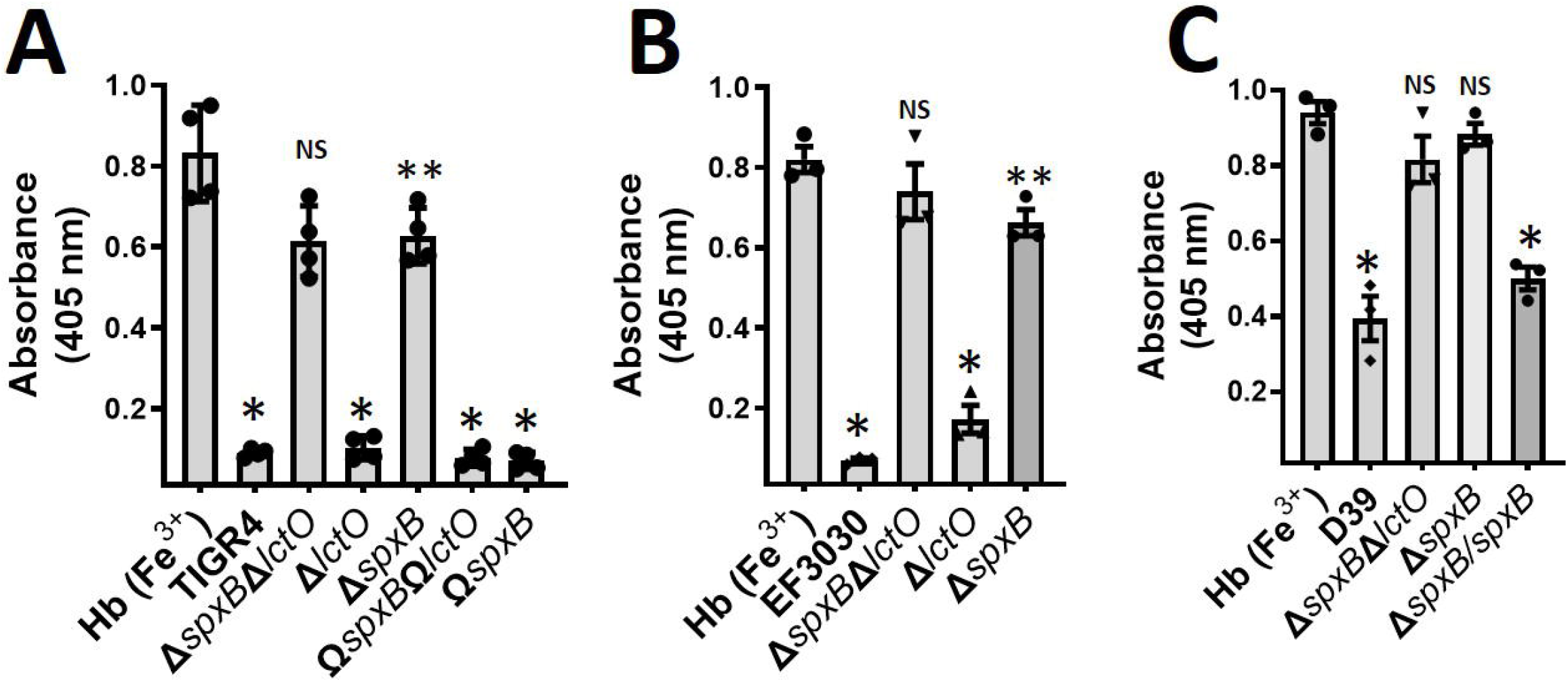
Oxidation of hemoglobin occurs through hydrogen peroxide produced through SpxB activity but not LctO. (A) TIGR4, TlGR4Δ*spxB*Δ*lctO*, TIGR4Δ*spxB*, TIGR4Δ*lctO*, TIGPAΩ*spxB*Ω*lctO*, or TIGR4Ω_spxB_ or (B) EF3030, EF3030Δ_spxB_Δ*lctO*, EF3030Δ_spxB_, or EF3030Δ*lctO*, or (C) D39, D39 Δ*spxB*Δ*lctO*, D39 Δ*spxB*, or D39 Δ*spxB/spxB* were inoculated in THY containing met-hemoglobin [Hb (Fe^3+^)] and cultures were incubated for 4 h at 37°C in a 5% CO_2_ atmosphere. Supernatants were collected and the spectra were obtained using a spectrophotometer Omega BMG LabTech. The absorbance at 405 nm obtained from each assessed strain, was utilized to construct the graphics. Error bars in all panels represent the standard errors of the means calculated using data from at least three independent experiments performed with two technical replicates each. One-way ANOVA with Dunnet’s test for multiple comparison was performed; NS=not significant compared to THY-Hb (Fe^3+^), \**p*<0.0001 compared with untreated THY-Hb (Fe^3+^), or Δ*spxB*Δ*lctO*, or Δ*spxB*; \*\**p*<0.003 compared with untreated THY-Hb (Fe^3+^).

### Spn-produced H_2_O_2_ degrades heme-Fe^3+^

Heme degradation occurs through enzymatic reactions or by nonselective destruction of heme double bonds caused by reactive oxygen species, such as H_2_O_2_ (35, 56). To investigate whether Spn-H_2_O_2_ caused degradation of heme, THY-Hb medium was inoculated with TIGR4, EF3030, their hydrogen peroxide mutant derivatives, or complemented strains and heme was assessed in supernatants using an in-gel heme assay (57, 58). Signal of heme, heme-bound to hemoglobin monomers or dimers, was detected in uninfected THY-Hb cultures incubated for 6 h (Fig. 4A). Heme signal in the supernatant of THY-Hb cultures incubated with TIGR4 for 2 h was similar to that of the uninoculated control but it decreased after 4 h post-inoculation and was not detected 6 h post-inoculation (Fig. 4A). In THY-Hb cultures of TIGR4, TIGR4Δ*lctO*, TIGPAΩ*spxB*Ω*lctO*, TIGP4Δ*spxB*Ω*spxB*, EF3030, or EF3030Δ*lctO* the signal of heme had already disappeared after 6 h of incubation (Fig. 4B and 4C). In contrast, in THY-Hb cultures of TIGPAΔ*spxB*Δ*lctO*, TIGR4Δ*spxB*, EF3030Δ*spxB*Δ*lctO*, or EF3030Δ*spxB*, signal of free heme, and heme-bound to hemoglobin was detected, thus confirming that degradation of heme had occurred in these cultures through hydrogen peroxide produced by SpxB.

**Fig. 4.**
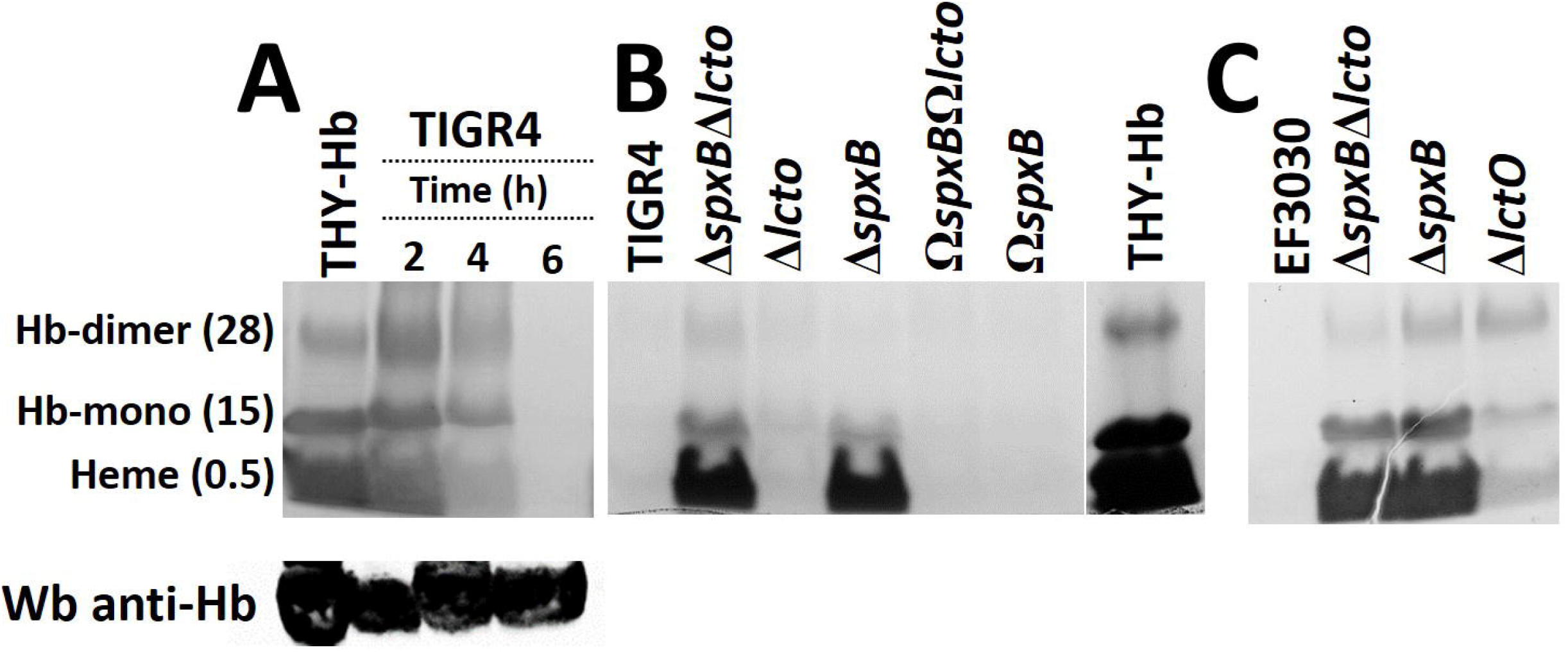
Oxidation of human hemoglobin by Spn-H_2_O_2_ leads to heme degradation. (A-C) The samples were stained by an in-gel staining method. (A) TIGR4 wt was inoculated in THY supplemented with met-hemoglobin (THY-Hb) and incubated for 2, 4 or 6 h at 37°C in a 5% CO_2_ atmosphere. Supernatant was harvested and run in a 12% polyacrylamide gel under non-denaturing conditions and heme was stained by an in-gel heme staining procedure. Free heme, heme-bound to hemoglobin monomer (Hb-monomer) or to hemoglobin dimers (Hb-dimer), and the observed molecular size in kDa, are indicated. In the bottom panel, supernatants were Western blotted using a polyclonal anti-hemoglobin antibody. (B) TIGR4, TIGR4Δ*spxB*Δ*lctO*, TIGR4Δ*lctO*, TIGR4Δ*spxB*, TIGR4Ω*spxB*, TIGR4Ω*spxB*Ω*lctO*, or (C) EF3030 wt strains or EF3030Δ*spxB*Δ*lctO*, EF3030Δ*spxB*, or EF3030Δ*lctO*, was inoculated into THY-Hb and incubated for six hours. Supernatants were harvested and analyzed as in (A).

As a loading control and to additionally investigate if hemoglobin had been degraded, supernatants were probed by Western blot with a polyclonal anti-hemoglobin antibody. Our results revealed a similar concentration of hemoglobin in those supernatants, confirming that the apo-protein was not degraded after the 2, 4 or 6 h incubation period with Spn (Fig. 4A, bottom panel). As expected since heme was degraded by Spn-H_2_O_2_, the intracellular concentration of iron was 79.19, 18.21, or 108.79 μM of Fe/μg of protein in TIGR4, TIGPAΔ*spxB*Δ*lctO* or TIGPAΩ*spxB*Ω*lctO*, respectively, confirming that iron was released and taken by wt and complemented strain. Altogether, these experiments demonstrated that Spn-H_2_O_2_ degrades free-heme and heme bound to hemoglobin.

### Streptococci that produces hydrogen peroxide, but not other respiratory bacteria, causes the degradation of heme

Non-encapsulated Spn (NESpn) and other alpha-hemolytic streptococci produce hydrogen peroxide (59, 60). We therefore assessed a collection of NESpn and oral streptococci for heme degradation. We included S. *mutans* as an additional control since strains lack production of H_2_O_2_ (61, 62). As shown in Fig. 5, degradation of heme-Fe^3+^was observed in THY-Hb cultures of NESpn strains MNZ41, MNZ67 and S. *salivarius, S. mitis, S. pseudopneumonanie* and S. *oralis* but not in cultures of S. *mutans* (Fig. 5A). We also tested cultures of other bacterial species that cohabit the upper airways along with oral streptococci, such as *Haemophilus influenzae, Staphylococcus aureus* and assessed cultures of *Klebsiella pneumoniae*. The incubation of THY-Hb with any of these bacterial species did not cause heme degradation (Fig. 5A). Streptococcal species that caused heme degradation produced hydrogen peroxide, while those unable to affect the heme-hemoglobin Soret curve did not (Fig. 5B). Altogether, these experiments demonstrated that Spn and other alpha hemolytic streptococci that produce abundant hydrogen peroxide, oxidizes hemoglobin causing the degradation of heme.

**Fig. 5.**
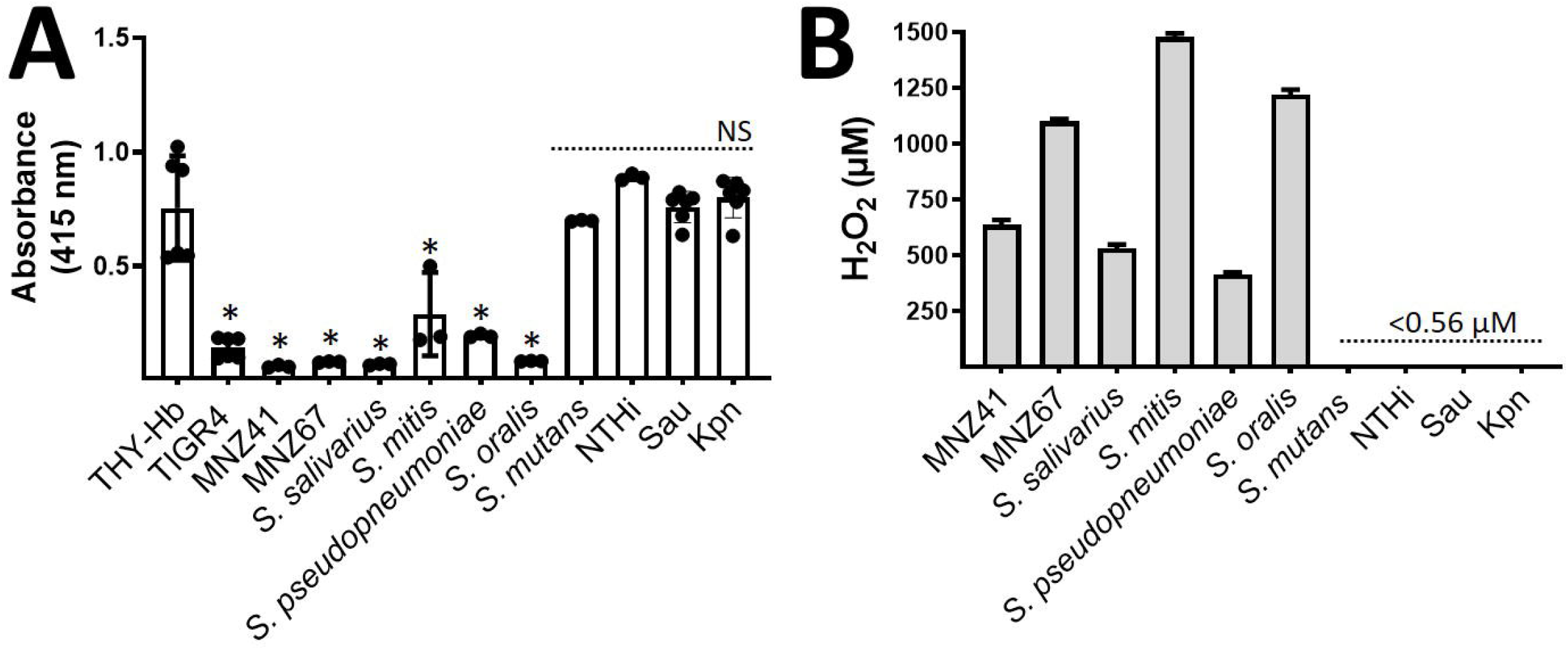
Oxidation of hemoglobin occurs in cultures of oral Streptococci that produce hydrogen peroxide. THY supplemented with met-hemoglobin (THY-Hb) was left uninoculated or inoculated with the indicated strain and incubated for 4 h at 37°C in a 5% CO_2_ atmosphere. Supernatants were collected and the spectra were obtained using a spectrophotometer Omega BMG LabTech. The absorbance obtained at 405 nm for each assessed strain, was utilized to construct the graphic. Error bars represent the standard errors of the means calculated using data from at least three independent experiments performed with two technical replicates. One-way ANOVA with Dunnet’s test for multiple comparison was performed, \**p*<0.05, or NS=not significant, compared with untreated THY-Hb. (B) Hydrogen peroxide was quantified from 4 h culture supernatants of each strain and using the Amplex Red assay. Error bars represent the standard errors of the means calculated using data from two independent experiments performed with two technical replicates. Limit of detection=0.56 μM.

### Heme is oxidized and degraded in an *ex-vivo* model of pneumococcal bacteremia

During the course of pneumococcal bacteremia erythrocytes are lysed by the pneumolysin (Ply), released through autolysis (63). Ply is also located on the bacterial membrane where it lyses red blood cells through contact (19, 20). Abundant Spn-H_2_O_2_ is produce as early as 2 h post-inoculation (Fig. 2D) thereby we sought to assess whether oxidation of hemoglobin and heme degradation occur in an *e×-vivo* model of pneumococcal bacteremia (Fig. 6A). To assess this, we supplemented THY broth with a suspension of red blood cells (THY-RBC), and this medium was infected with TIGR4, TIGR4Δ*ply*, or TIGR4Δ*spxB*Δ*lctO*. The oxidation and concentration of hemoglobin as well as heme degradation were investigated.

**Fig. 6.**
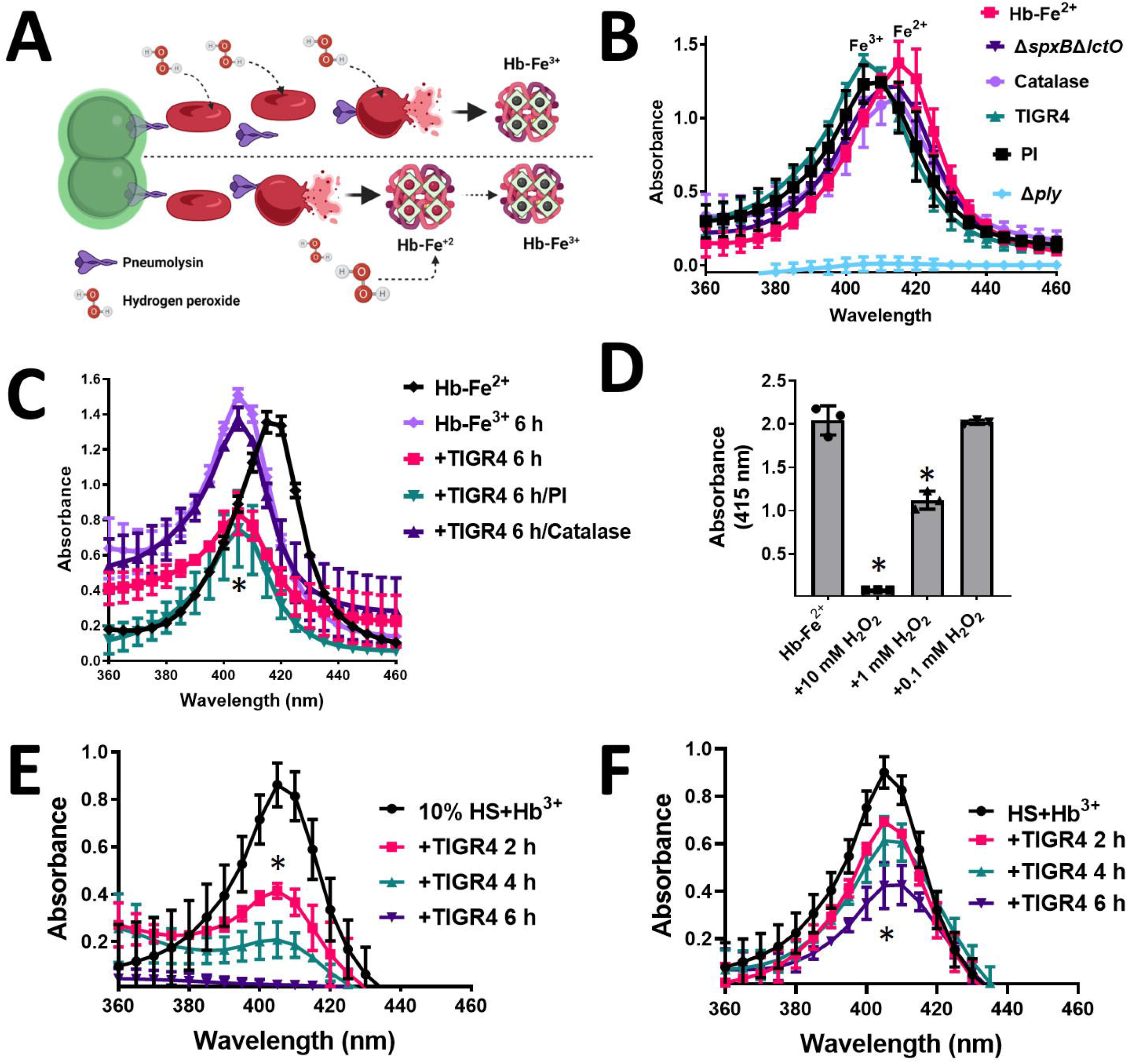
Hemoglobin is oxidized by Spn-H_2_O_2_ in an ex *vivo* bacteremia model with erythrocytes. (A) Model of *ex vivo* pneumococcal bacteremia showing pneumococci with pneumolysin (Ply) located in the membrane; Ply also gets released through autolysis. Membrane-permeable hydrogen peroxide (H_2_O_2_) is produced as a by-product of the oxidation of pyruvate. Bottom panel, oxy-hemoglobin (Hb-Fe^2+^) is then release from erythrocytes by Ply-induced erythrocyte lyses and once released, oxyhemoglobin gets oxidized by Spn-H_2_O_2_ to met-hemoglobin (Hb-Fe^3+^). Alternatively, top panel, Spn-H_2_O_2_ permeates inside erythrocytes causing the oxidation of hemoglobin that is then released from lysed erythrocytes as met-hemoglobin (Hb-Fe^3+^). (B and C) A suspension (2%) of sheep erythrocytes in THY was inoculated with TIGR4, TIGR4 supplemented with catalase, TIGR4 supplemented with protease inhibitors (PI), TIGR4Δ*spxB*Δ*lctO*, or TIGR4Δ*ply* and the inoculated cultures were incubated for 30 min (B) or 6 h (C) at 37°C in a 5% CO_2_ atmosphere. As a control, erythrocytes were lysed with saponin and oxy-hemoglobin (THY-Hb^2+^), that was released into the supernatant, was considered as maximum hemoglobin release. (D) Oxy-hemoglobin was left untreated (Hb-Fe^2+^) or added with added with H_2_O_2_ and incubated for 6 h.(E and F) Human serum was supplemented with 10 μM methemoglobin (HS+Hb^3+^) or serum was used to supplement THY to a final 10% (v/v) and added with 10 μM met-hemoglobin (10% HS+Hb^3+^). These media were inoculated with TIGR4 and incubated for 2, 4 or 6 h. Supernatants from experiments in panels (B-F) were collected and the spectra were obtained using a spectrophotometer Omega BMG LabTech. In panels B through F, error bars represent the standard errors of the means calculated using data from at least three independent experiments. One-way ANOVA with Dunnet’s test for multiple comparison was performed, \**p*<0.05, or NS=not significant, compared with (B and C) TIGR4/catalase, (D) untreated Hb-Fe^2+^, (E) untreated 10% HS+Hb^3+^, or (F) untreated HS+Hb^3++^.

As expected, the Soret curve was not observed in supernatants from non-lysed THY-RBC control and it was also absent in THY-RBC supernatants infected with TIGR4Δ*ply*, because it does not lyse erythrocytes (Fig. 6B). Given that RBCs contains oxy-hemoglobin (Hb-Fe^2+^), the Soret curve of the positive control, where RBC had been lysed with saponin, peaked at 415 nm with an average absorbance of 1.38 (Fig. 6B). The concentration of cell-free hemoglobin in this control supernatant was 17.35 μM as measured by the Quantichrom assay (Table 1). The Soret curve from THY-RBC cultures infected with TIGR4 peaked at 405 nm with an average absorbance of 1.39 (Fig. 6B) and concentration of hemoglobin of 12.8 μM (Table 1). These results indicate that maximum TIGR4-induced release of hemoglobin is achieved within 30 min of incubation, compared with the saponin-induced released control, and that the oxidation of heme-Fe^2+^ to heme-Fe^3+^ had already occurred (Fig. 6B). Similar to the wt and because TIGR4Δ*spxB*Δ*lctO* produces Ply (21, 23), a Soret curve was observed in THY-RBC cultures infected with this hydrogen peroxide deficient mutant as soon as 30 min post-inoculation (Fig. 6B). This Soret curve peaked at 415 nm with an average absorbance of 1.21 and a concentration of hemoglobin of 16.4 μM (Table 1), indicating that most hemoglobin had been released but, unlike the wt strain, oxidation of heme-Fe^2+^ did not occur because of the lack of hydrogen peroxide. Accordingly, oxidation of hemoglobin did not occur in TIGR4 cultures treated with catalase but it was oxidized to met-hemoglobin in TIGR4 cultures treated with protease inhibitors (Fig. 6B).

**Table 1.**
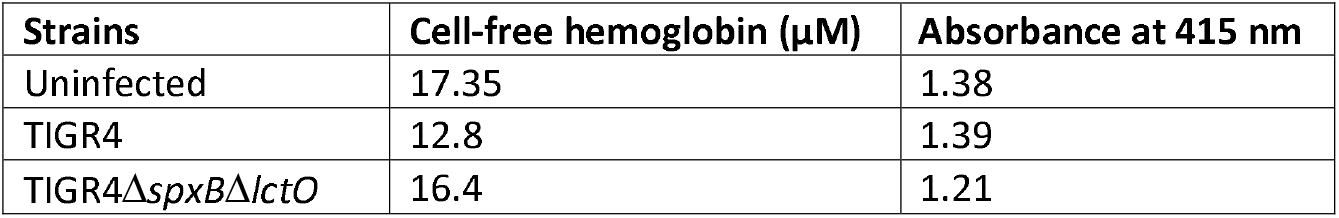
Quantification of cell-free hemoglobin and the heme-hemoglobin Soret curve.

After 6 h of incubation, autoxidation of uninoculated, saponin-released hemoglobin in THY-RBC cultures (Fig. 6C, Hb-Fe^3+^) and those infected with TIGR4 treated with catalase (Fig. 6C) were oxidized. THY-RBC medium infected with TIGR4 had already flattened the heme-Fe^3+^ curve and therefore heme had been degraded but not in those of TIGR4 treated with catalase (Fig. 6C). Degradation of heme was not inhibited by supplementing THY-RBC before the infection with protease inhibitors (Fig. 6C). We next treated saponin-released oxy-hemoglobin with increasing amounts of H_2_O_2_ and the oxidation to met-hemoglobin and then degradation of heme was recapitulated (Fig. 6D). Altogether, our results indicate that oxidation of hemoglobin and degradation of heme by Spn-H_2_O_2_ occur during pneumococcal bacteremia.

To further mimic bacteremia, we added THY with 10% of human serum and supplemented this broth culture with 10 μM met-hemoglobin. In addition, non-diluted human serum was supplemented with 10 μM met-hemoglobin. These hemoglobin-containing serum cultures were infected with TIGR4 and incubated at different times. A time course degradation of met-hemoglobin was observed in both culture conditions (Fig. 6E and 6F), confirming that hemoglobin is oxidized and degraded when Spn growths in human serum.

Because the release of hemoglobin and its oxidation was observed early during the simulated bacteremia (i.e., 30 min), to gain insights into the mechanism leading to hemoglobin oxidation and heme degradation, we investigated expression of *ply, spxB*, and *lctO* during incubation of Spn in THY-RBC, THY-Hb or in THY supplemented with heme. Compared to THY control cultures, the expression of *ply, spxB*, and *lctO* did not significantly change when TIGR4 grew in broth cultures added with RBCs or hemoglobin. However, transcription of *spxB* was upregulated when TIGR4 was incubated in THY containing heme, whereas the transcription *lctO* and *ply* was downregulated (Fig. 7). Together, these experiments indicate that under these culture conditions heme was the inducer for Spn-H_2_O_2_ production.

**Fig. 7.**
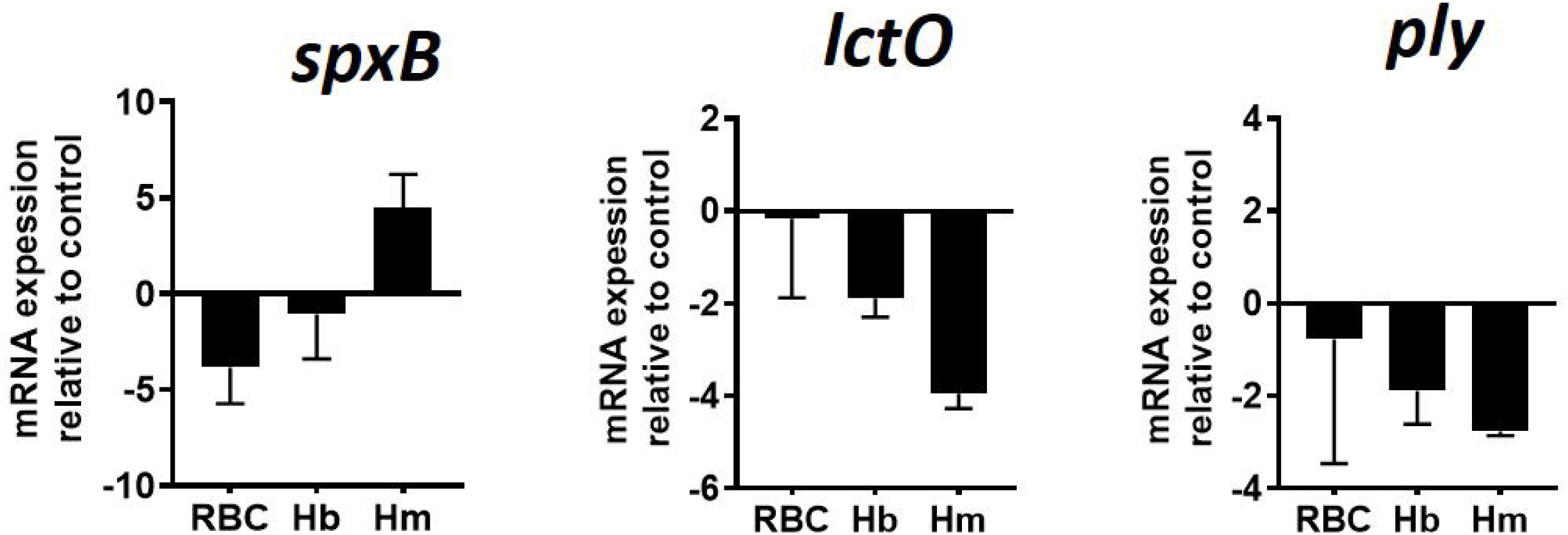
Regulation of *ply*, *spxB* and *lctO* genes by incubation with erythrocytes, hemoglobin, or heme. TIGR4 wt was inoculated in THY broth alone, in THY broth containing 1% sheep erythrocytes, THY with 10 μM hemoglobin (Hb) or THY with 5 μM heme. Cultures were incubated for 2 h at 37°C in a 5% CO_2_ atmosphere, bacteria were then harvested, and RNA was extracted from these cultures and utilized as template in qRT-PCR reactions with primers that amplified the *ply, spxB*, or *lctO* gene. Average *C_T_* values were normalized to the 16S rRNA gene, and the fold differences, relative to THY control, were calculated using the comparative *C_T_* method (2^-ΔΔ*C*^_*T*_)(78).

## Discussion

We have demonstrated in this study that H_2_O_2_ produced by the pneumococcus, Spn-H_2_O_2_, and other alpha-hemolytic Streptococci, caused the oxidation of hemoglobin to met-hemoglobin with a subsequent degradation of heme. The current study also demonstrated that the oxidation of hemoglobin and heme degradation occurred in a simulated bacteremia infection suggesting that these oxidative reactions may occur during pneumococcal disease.

*In vitro* oxidation of hemoglobin with hydrogen peroxide produces ferryl hemoglobin [Hb-Fe^4+^], a highly reactive form of hemoglobin that gets reduced to met-hemoglobin (64). With an increased presence of hydrogen peroxide molecules, met-hemoglobin gets oxidized back to Hb-Fe^4+^, which in turn causes heme degradation (65). Iron stored in heme-hemoglobin is poorly reactive but once freed up, as a consequence of heme degradation, free iron can react with hydrogen peroxide to produce another highly reactive hydroxyl radical (66, 67). Indeed, we have demonstrated that free iron present in THY medium is enough to generate hydroxyl radicals (^•^OH) (40), killing S. *aureus* strains rapidly (39, 68, 69). The increase of iron in the culture medium due to heme degradation should result in an increased generation of ^•^OH radicals. The potential presence of two highly reactive radicals, Hb-Fe^4+^ and ^•^OH, for the cytotoxicity seen during pneumococcal disease is being investigated in our laboratories.

It has been investigated that nearly 3% of oxy-hemoglobin is auto-oxidized inside erythrocytes, and this auto-oxidation is caused by 2×10^-7^ mM of hydrogen peroxide generated in the cytosol of red blood cells (70). Although auto-oxidation by CO_2_ occurred in our *in vitro* system 4 h post-inoculation (i.e., auto-oxidation was not observed when incubated at 37°C with no CO_2_), Spn cultures produce 10^7^-fold more (~1 mM) hydrogen peroxide within a few hours of incubation and caused the rapid oxidation of hemoglobin leading to the release of heme, a toxic molecule, with its subsequent degradation (65). We demonstrated that heme degradation caused an increased on free-iron in the medium that was taken by pneumococci. These novel observations might have important implications for pathogenesis of hydrogen peroxide-producing Streptococci but also for the symbiosis observed in bacterial communities where some species can benefit from heme and iron release by hydrogen peroxide-producing bacteria.

In children with SCD, hemoglobin (Hb)S is oxidized to Hb-Fe^4+^ then reduced to methemoglobin and resulting in an acute hemolytic vascular inflammatory process causing acute lung injury (30, 50). Both Hb-Fe^4+^ and met-hemoglobin induce a drop of mitochondrial oxygen consumption and mitochondrial membrane potential in epithelial lung cells (36). Acute chest syndrome is an important cause of hospitalization and mortality in children with SCD and Spn is a leading etiologic agent of acute chest syndrome. While the precise role of Spn is still unclear, SCD children are ~100-fold more susceptible to pneumococcal infection (71). Studies of the oxidation of hemoglobin by Spn-H_2_O_2_, however, have been neglected. We recently demonstrated that the growth of planktonic Spn, and formation of pneumococcal biofilms, is enhanced by supplementing the culture medium with human hemoglobin (72, 73). Besides iron, other nutrients may become accessible to pneumococcus, since the oxidation of hemoglobin by hydrogen peroxide releases globin-derived peptides (29), but also a striking transcriptome remodeling was identified that included upregulated transcription of genes encoding transporters of glyco-conjugated molecules (72, 73).

Besides nutrients, oxidative reactions may lead to unintended consequences. During pneumococcal pneumonia the pulmonary parenchyma presents with hemorrhage, inflammatory congestion, hepatization, suppurative infiltration, and lung parenchymal injury (4, 9–13). Whereas there are a number of virulence factors implicated in the pathophysiology of lung infection (63), the oxidation of hemoglobin through Spn-H_2_O_2_ may also be an important contributor to pneumococcal disease. For example, mice intranasally inoculated with Spn develop lung hemorrhage, at necropsy, and histological analysis reveals lung consolidation associated with alveolar septal edema and pleomorphic alveolar, interstitial, and perivascular cellular inflammation (74). Spn mutants in the *spxB* gene whether with a pleotropic capsule defect or not, were attenuated for virulence in mouse models of pneumococcal disease (46, 47, 75).

There are other important implications as a result of the oxidation of hemoglobin by hydrogen peroxide produced by streptococci. Heme release and/or heme degradation through, hydrogen peroxide, may be beneficial to other bacteria which do not synthetize large amounts of this pro-oxidant but that require iron for their metabolism and pathogenesis. For instance, it has been recently demonstrated that dental plaque bacteria such as S. *gordonii*, which produces hydrogen peroxide, oxidizes hemoglobin to release heme, and this free-heme facilitates the colonization of the dental plaque by a Gram-negative strain *Porphyromonas gingivalis* (49). *P. gingivalis* does not produce siderophores but it has a sophisticated mechanism for heme-iron acquisition (76). Conversion of oxy-hemoglobin to met-hemoglobin, by hydrogen peroxide producing bacteria, allows the release of heme that can be taken by these microorganisms.

In summary, we demonstrated that Spn and other hydrogen peroxide-producing Streptococci, oxidizes hemoglobin from human and other species, releasing and degrading heme. These oxidative reactions occurred early during the lag phase of growth indicating that metabolic adaptation for growth in the presence of hemoglobin stimulated production of Spn-H_2_O_2_. This upregulated metabolic adaption was the result in part of the release of heme since gene expression studies demonstrated an upregulation of *spxB* transcription but not that of *lctO* or the pneumolysin gene *ply*, under the culture conditions tested. Noteworthy, incubation with erythrocytes or hemoglobin did not induced a regulation of *spxB* in the current study. We have observed a potential toxic radical produced during the oxidation of hemoglobin, spanning the incubation time utilized in gene expression studies, that may explain the observed downregulation of *spxB* (Scasny et al, unpublished observations). Efforts are in place to assess the specific role for these oxidative reactions for pathogenesis and bacterial symbiosis.

## Material and Methods

### Bacterial strains, material, and growth conditions

All Spn wild type strains and mutant derivatives used in this study are listed in Table 2. Bacterial stocks were prepared in medium containing skim milk-tryptone-glucose-glycerin (STGG) and stored at −80°C (77). Pneumococci were rotinously cultivated at 37°C in a 5% CO_2_ and ~20% O_2_ atmosphere on blood agar plate (BAP). Non-typeable *Haemophilus influenzae* (NTHI) was grown on chocolate agar plate (CAP), which was prepared with heat-lysed horse blood, a source of hemin and NAD. *Klebsiella pneumoniae* and *Staphylococcus aureus* were cultivated on tryptic soy agar (TSA). Experiments were performed using Todd-Hewitt-broth with 0.5% yeast extract (THY), THY added with 10 μM human met-hemoglobin (Sigma-Aldrich), THY added with 5 μM heme (Sigma-Aldrich), THY added with sheep erythrocytes (Quad Five), and horse erythrocytes (Lampire). In experiments inoculated with NTHI, THY medium was supplemented with hemin (10 μg/ml), and NAD (2 μg/ml). Brain heart infusion (BHI) broth, Bacto agar, Bacto Tryptic Soy Broth, Bact Yeast Extract and Bacto Todd Hewitt Broth were purchased from Becton, Dickinson and Company. Antibiotics utilized were gentamycin, erythromycin, spectinomycin, chloramphenicol, and were all sourced from Sigma-Aldrich. Hydrogen peroxide (Fisher Scientific), Nicotinamide adenine dinucleotide (NAD(+)) (Sigma-Aldrich), Catalase (Sigma-Aldrich), Complete™, Mini Protease Inhibitor Cocktail (Roche), FeCl_3_·6H_2_O and potassium hexacyanoferrate (III) (Sigma-Aldrich), o-dianisidine (Alfa Aesar) /sodium acetate (Sigma-Aldrich), ethanol and methanol (Fisher Scientific). Human serum was purchased from MP Biomedicals.

**Table 2.**
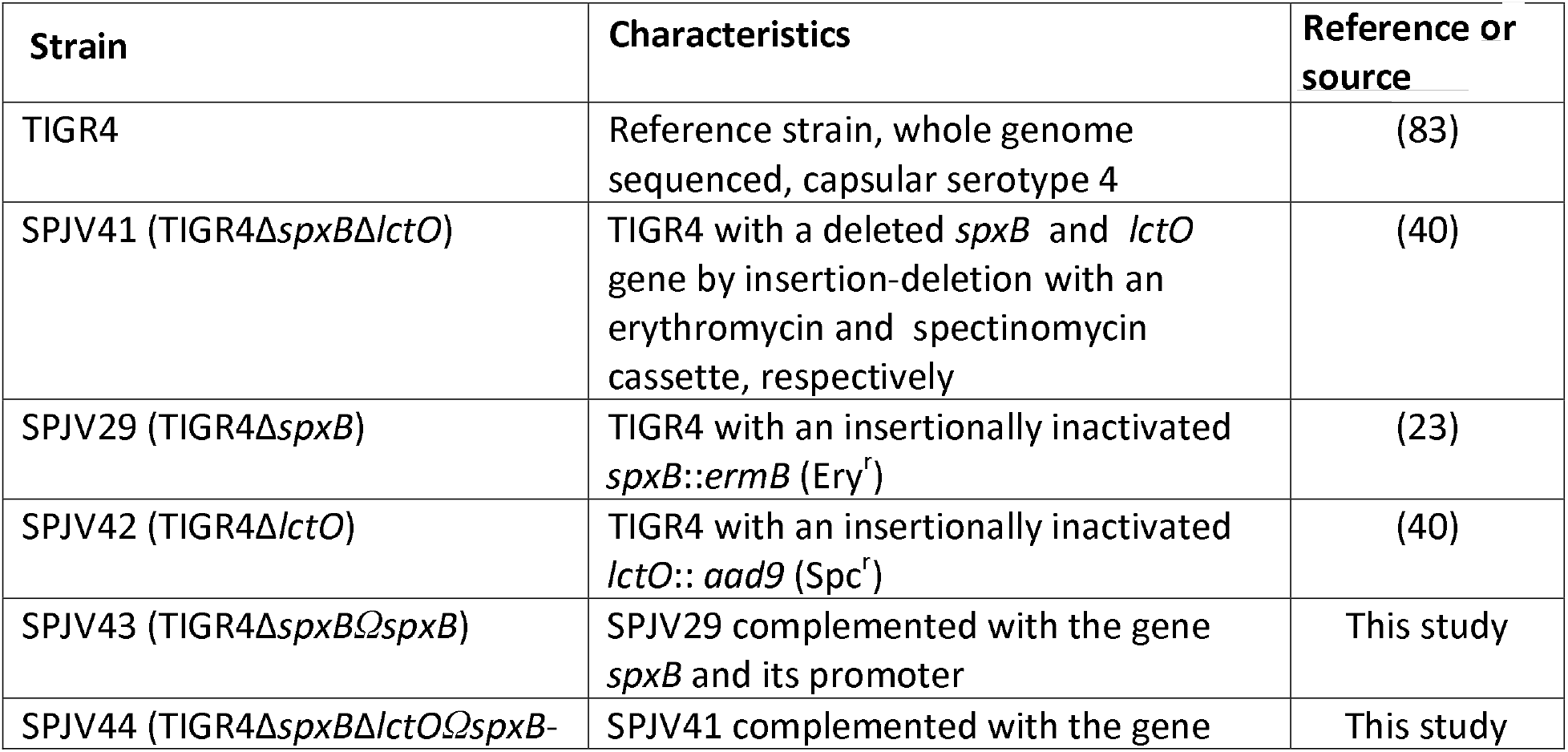
Strains utilized in this study.

### Preparation of inoculum for experiments

The inoculum was prepared as previously described (40). Briefly, bacteria were inoculated on blood agar plates (BAP) and incubated overnight at 37°C in a 5% CO_2_ atmosphere. Bacteria were then harvested from plates by PBS washes and this suspension was used to inoculate THY with or without additives, that was brought to a final OD_600_ of ~0.1. This suspension contained ~5.15×10^8^ cfu/ml as confirmed by dilution and platting of aliquots of the suspension.

### Bacterial broth culture medium containing oxy-hemoglobin

Sheep or horse erythrocytes were washed three times with sterile PBS (pH=7.4) at 300⍰×⍰*g* for 5⍰min in a refrigerated centrifuge (Eppendorf). These washed erythrocytes suspension was lysed with 0.1% saponin and the concentration of oxy-hemoglobin in this preparation was determined using the QuantiChrom™ Hemoglobin Assay Kit (BioAssay Systems); oxy-hemoglobin was then added to THY broth to a final concentration of ~10 μM. The concentration, and redox state of oxy-hemoglobin was verified by spectroscopy before each experiment. In some experiments, THY containing oxy-hemoglobin was supplemented with an excess of catalase to a final concentration of 400 U/ml, or with 1X complete protease inhibitors cocktail (Roche).

### Studies of the oxidation of oxy-hemoglobin, met-hemoglobin and heme degradation by spectroscopy

Experiments were conducted using six-well microplates (Genesee Scientific) added with THY containing a 2% suspension of washed sheep erythrocytes, 10 μM of oxy-hemoglobin, or 10 μM of met-hemoglobin. In some experiments human serum was supplemented with 10 μM of met-hemoglobin, or THY was supplemented with human serum to a final 10% concentration (v/v) and added with 10 μM of met-hemoglobin. These media were inoculated with Spn or the other bacterial species listed in each experiment, and incubated for the indicated time at 37°C in a 5% CO_2_ and ~20% O_2_ atmosphere. At the of the incubation time, supernatants containing planktonic bacteria were removed from cultures and bacteria were pelleted down by centrifugation at 13,000 rpm x 5 min in a refrigerated micro-centrifuge (Eppendorf). In another set of experiments, THY containing 10 μM of human met-hemoglobin was treated with different concentrations of hydrogen peroxide for 6 h. The oxidation of oxy-hemoglobin to met-hemoglobin and further oxidation and degradation of met-hemoglobin was determined by analyzing the spectra spanning 200 nm through 1000 nm in bacteria-free supernatants, using a spectrophotometer Omega BMG LabTech (ThermoFisher).

### Quantification of hydrogen peroxide production

Hydrogen peroxide production was quantified from bacterial cultures inoculated in six-well microplates (Genesee Scientific) that had been added with THY. These cultures were incubated at 37°C in a 5%CO_2_ and ~20% O_2_ atmosphere for the indicated time. At the end of the incubation, the culture supernatant was collected, centrifuged at 4°C for 5 min at 15,000 rpm and then the supernatant was transferred to a new tube and kept in ice for no more than 30 min. Some supernatants were filtered sterilized using a 0.4 μm syringe filter (Fisher scientific). Collected supernatants were diluted with 1× reaction buffer of the Amplex Red H_2_O_2_ assay kit (Molecular Probes), and hydrogen peroxide levels were quantified according to the manufacturer’s instructions using a spectrophotometer Omega BMG LabTech (ThermoFisher).

### Qualitative detection of hydrogen peroxide production by *S. pneumoniae* strains

Qualitative detection of H_2_O_2_ production by Spn was performed using the Prussian blue agar assay, which limit of detection is 2.5 nm H_2_O_2_ (55). Prussian blue agar plates were made with 1.0 g/liter of FeCl_3_·6H_2_O, 1.0 g/liter of potassium hexacyanoferrate III (K_3_[Fe(CN)_6_]) also known as Prussian blue, 37 g/liter of dehydrated BHI broth, and 15 g/liter of agar. Prior to experiments, detection of H_2_O_2_ was assessed by dropping 10 μl of a 1 mM H_2_O_2_ solution that reduced K_3_[Fe(CN)_6_] and causing its precipitation and then a blue halo within minutes. PBS was used as a negative control. Spn strains or culture supernatants (10 μl) obtained as described above, were inoculated, or spotted, respectively, onto Prussian Blue agar plates. These plates were incubated at 37 °C in a 5% CO_2_ and ~20% O_2_ atmosphere overnight (Spn) or for 2 h (supernatants) after which plates were photographed with a Canon Rebel EOS T5 camera system and digital pictures were analyzed.

### Western blotting

Bacteria were grown in THY with 10 μM hemoglobin at 37°C for 2 h, 4 h, and 6 h. After incubation the cells were harvested by centrifuge at 15,000 rpm for 5 min and the supernatant were transferred to new tube. The collected samples were combined with 4x reducing sample buffer and boiled 5 minutes. The mixture was loaded into 10 well 4-12% Mini-PROTEAN^®^ TGX™ precast gel (Biorad), and run 2 hours at 90 V in running buffer. Gels were transferred to nitrocellulose membranes in transfer buffer supplemented with 20% ethanol with Trans-Blot Turbo Transfer System. Membranes were blocked 1 h at room temperature in 5% non-fat dry milk in TBS supplemented with 0.1% Tween-20 (TBST). Then a membrane was incubated with the Hb polyclonal antibody (Invitrogen) at 1:1000 dilution in 5% non-fat dry milk in TBST overnight at 4°C. Next day, the blot was washed 3x 5 min with TBST, and incubated with donkey anti-goat IgG-HRP (Santa Cruz) secondary antibody 1:5000 in 5% non-fat dry milk in TBST 1 h at room temperature. Blots were washed 3x 5 min in TBST and 2mL each SuperSignal Substrate (Thermo Fisher Scientific) reagent was added to blot and incubated for 5 min at room temperature. Blots were imaged on a ChemiDoc MP (BioRad) using ImageLab 5.0 software, automatic exposure settings for Chemiluminescence, high specificity, optimizing for bright bands.

### In-gel heme staining

Bacteria were grown in THY with 10 μM hemoglobin at 37°C for 2, 4, or 6 h, after which supernatants were collected, spun down and combined with non-reducing loading buffer. The mixtures were loaded into a non-denaturing 12% Mini-PROTEAN® TGX™ precast gel (Biorad), and run for 2 h at 90 V in running buffer lacking SDS. The gel was then stained using a published protocol (35). Briefly, after electrophoresis gels were immersed into a methanol/sodium acetate (pH=5) solution and incubated at room temperature on rocking platform (VWR) at 1 speed for 2 min with constant shaking. After incubation, the gel was added with a solution of o-dianisidine/sodium acetate (pH=5)and incubated for 20 min in the dark. To visualize heme in the gel, 3%hydrogen peroxide was added, and the reaction was stopped by washing with distilled water. The gel was then photographed with a Canon Rebel EOS T5 camera system and digital pictures were analyzed.

### RNA extraction and qRT-PCR analysis

Bacteria were inoculated into THY, or THY containing 5 μM heme, 5 μM Hb, or a 1% suspension of RBC and incubated at 37°C for 2 h. After incubation the bacterial suspension was collected, and mixed with RNAprotect Reagent (Qiagen). Then cells were harvested and the total RNA was extracted using the RNeasy mini kit (Qiagen) following the manufacturer’s instruction. DNA was removed using the Turbo DNase-free kit (Life Technologies). The RNA concentration was obtained using a NanoDrop spectrophotometer (Thermo Fisher Scientific), and 200 ng of RNA were cDNA transcribed using the iScript™ cDNA Synthesis Kit (Biorad). Gene expression analysis was carried out using the PerfeCTa SYBR® Green SuperMix (Quantabio) and CFX96 Touch Real-Time PCR system (Biorad). Primers used for the qRT-PCR analysis are listed in Table 3. The following conditions were utilized: 1 cycle at 95°C for 3 min and 40 cycles of 95°C for 15 s, 60°C for 30 s and 72°C for 30 s. Melting curves were generated to confirm the absence of primer dimers. The relative quantitation of mRNA expression was normalized to the constitutive expression of the housekeeping 16S rRNA gene and calculated by the comparative CT (2^-ΔΔCT^) method (78).

**Table 3.**
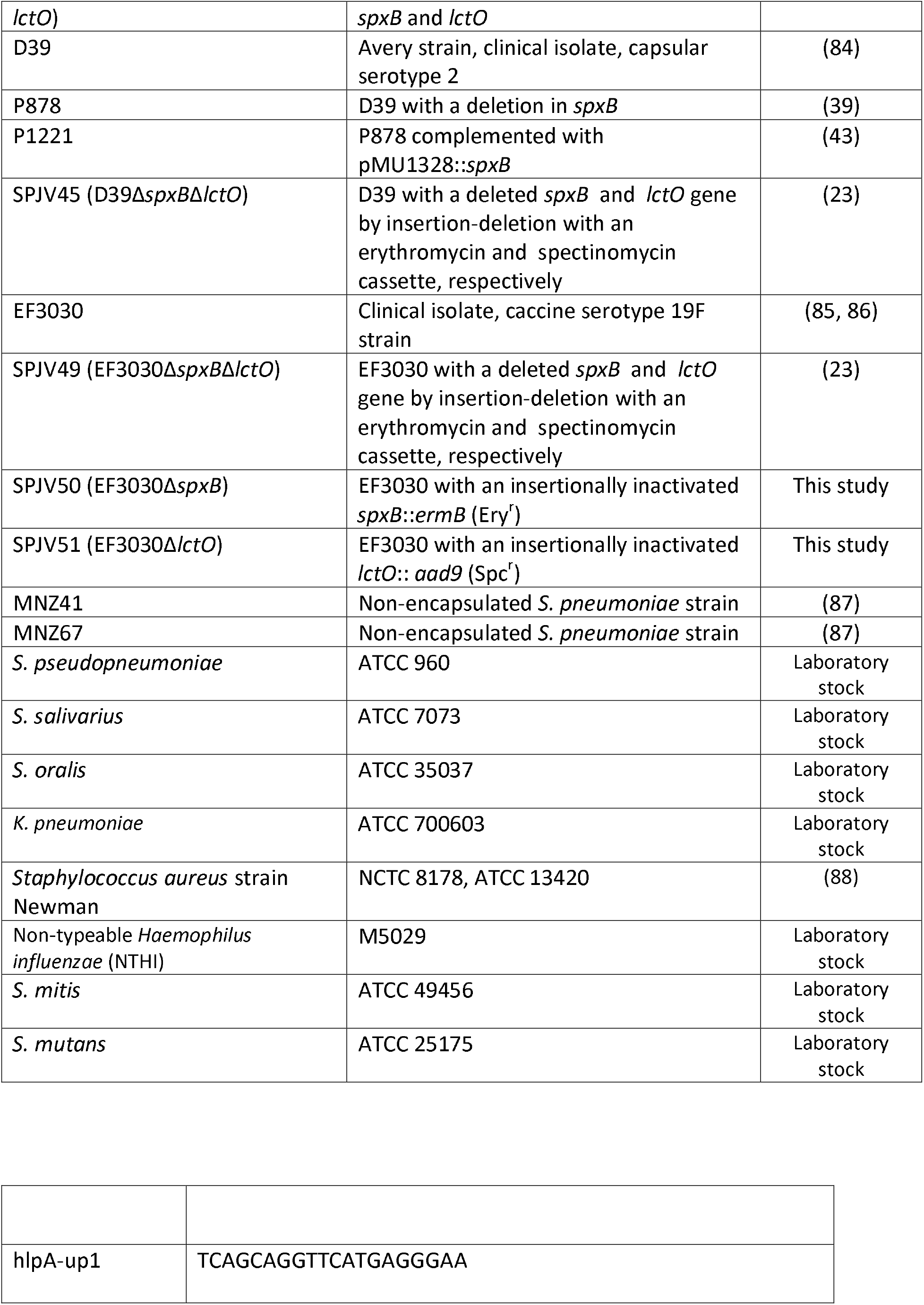
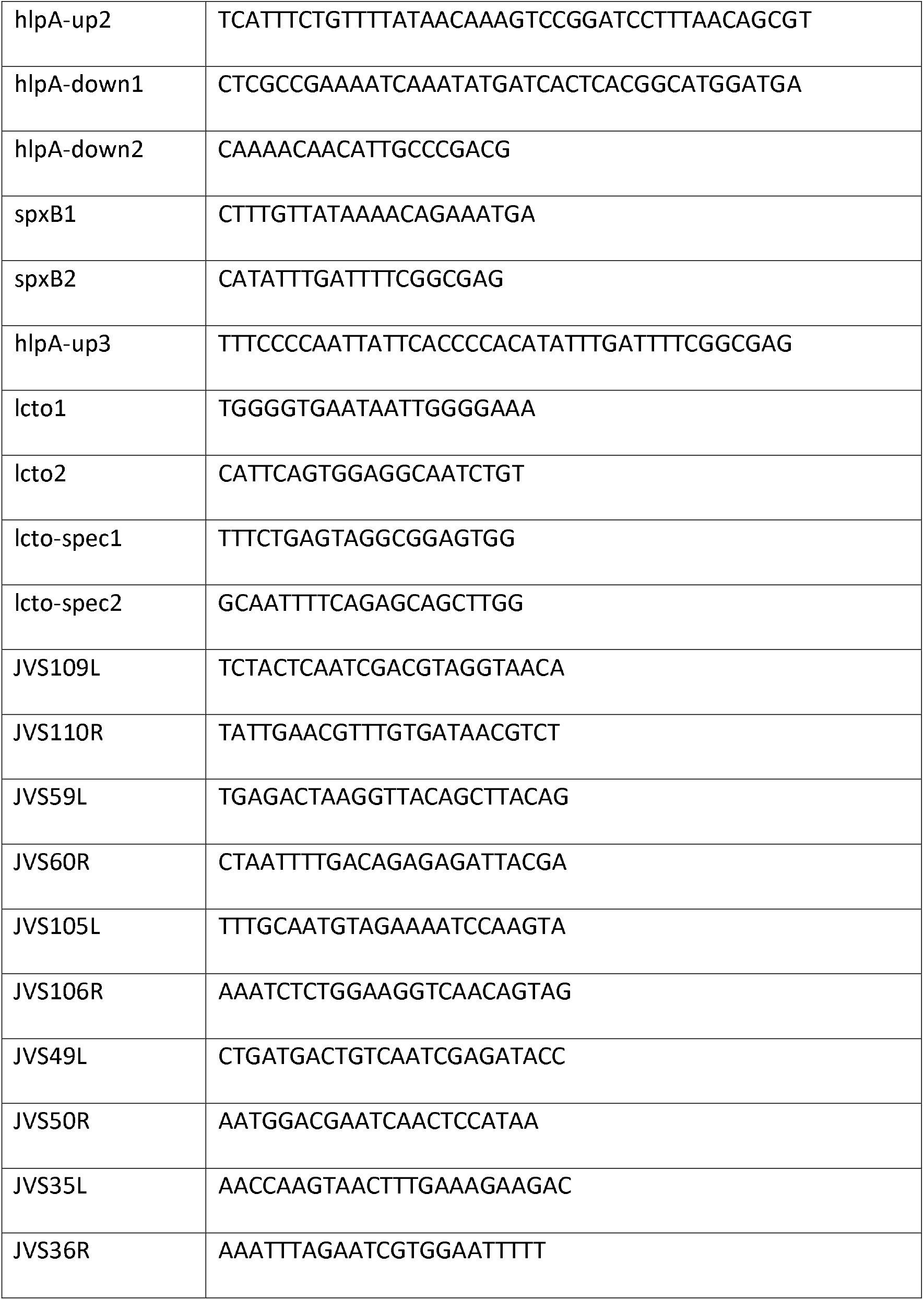
Oligo nucleotides designed and used in this study.

### Preparation of *lctO* mutation in Spn strains EF3030 and D39

To prepare an isogenic *lctO* mutant, we PCR-amplified, using primers lcto-spec1 and lcto-spec2 (Table 3), an insertionally inactivated *lctO*-spectinomycin fragment from strain TIGR4Δ*lctO* (40). This PCR product was purified using the QIAquick PCR purification kit (Qiagen) and then transformed into EF3030 or D39 using an standard transformation procedure (79). Transformants were harvested in BAP with spectinomycin (100⍰μg/ml) and the insertional inactivation of *lctO* was confirmed by PCR with primers lcto-spec1 / lcto-spec2 and sequencing.

### Construction of complemented strains

To complement *spxB*, or *spxB* and *lctO*, into TIGR4Δ*spxB* or TIGR4Δ*spxB*Δ*lctO*, respectively, we chose a ‘neutral’ chromosomal location within the *hplA* gene (SP1113) in which insertions do not affect virulence (52). A *spxB* complemented fragment was prepared by amplifying an upstream region carrying a fragment of the *hlpA* gene, using as a template DNA purified from TIGR4, and primers hlpA-up1 and hlpA-up2 (Table 3). We then generated a PCR fragment, using DNA purified from TIGR4 as a template, containing the *spxB* gene and its promotor with primers spxB1 and spxB2 (Table 3). A downstream fragment containing the *catP* gene from chloramphenicol resistance and a fragment of ORF SP1114 was amplified using as a template DNA from strain JWV500 using primers hplA-down1 and hlpA-down2. PCR products were purified using a QIAquick PCR purification kit (Qiagen) and ligated by splicing overlap extension PCR with primers hlpA-up1/hlpA-down2. The PCR-ligated product was verified in a 0.8% DNA gel, purified from the gel using QIAquick Gel Extraction Kit (Qiagen) and re-amplified by PCR with primers hlpA-up1 and hlpA-down2. This PCR product was purified as above and used to transform TIGR4Δ*spxB* using a standard procedure (79). Transformants were harvested in BAP plates containing chloramphenicol (5 μg/ml) and screened by PCR using primers spxB1/spxB2 to confirm the complementation of *spxB*.

To construct a *spxB* and *lctO* complemented strain, an upstream DNA fragment containing *hlpA* and *spxB* was amplified using DNA purified from TIGR4Δ*spxB*ΩspxB and primers hlpA1 and spxB2. Then, a DNA fragment was prepared by PCR amplification of the *lctO* gene and its putative promotor region using TIGR4 DNA as a template and primers lctO1 and lctO2. The downstream fragment containing the *catP* gene and a fragment of ORF SP1114 was used as prepared above. After purification with a QIAquick PCR purification kit (Qiagen), these three DNA fragments were ligated by splicing overlap extension PCR with primers hlpA-up1/hlpA-down2 and purified. This construct was re-amplified with primers hlpA-up1 and hlpA-down2, purified and transformed into TIGR4Δ*spxB*Δ*lctO*. Transformants were screened by PCR using primers hlpA-up1/hlpA-down2and selected completed clones were further confirmed by H_2_O_2_ production.

### Metal content analysis

Pneumococcal biofilms cultured in THY-Hb for 4 h were washed 3x with ice-cold PBS, resuspended in 1 ml PBS and sonicated at high intensity for 5 min in 30 s on 30 s off cycles using a Bioruptor (Diagenode, Denville, NJ, USA). Protein was determined in all samples by the Bradford method and a known mass of sample was mineralized in concentrated HNO_3_ trace metal grade (Thermo Fisher Scientific) as previously described (80). Iron in each sample was determined by atomic absorbance spectroscopy (AAS) using a flame 55 AA Absorption Spectrometer (Agilent, Santa Clara, CA, USA). Analytical grade standards for iron (Thermo Fischer Scientific) were diluted in 18ΩV water. The iron content on each sample was normalized to the initial mass of protein as previously reported for other metals (81, 82).

### Statistical analysis

We performed one-way analysis of variances (ANOVA) followed by Dunnet’s multiple comparison test, when more than two groups were compared or the Student *t* test to compare two groups, as indicated. All statistical analysis was performed using the software Graph Pad Prism (version 8.3.1).

## Acknowledgements

This study was supported in part by a grant from the National Institutes of Health (NIH; 5R21AI144571-03) to JEV and Wesleyan University institutional funds to TP-B. BA was supported by a Fulbright scholarship awarded by the US Department of State. FMF is supported by the Ronald E. McNair Postbaccalaureate Fellowship program. The content is solely the responsibility of the authors and does not necessarily represent the official view of the NIH or the US Department of State. The authors thank Jaime Carrazco-Carrillo from Wesleyan University for his assistance in some laboratory assays.

